# Accelerated DNA methylation aging and increased resilience in veterans: the biological cost for soldiering on

**DOI:** 10.1101/207688

**Authors:** Divya Mehta, Dagmar Bruenig, Bruce Lawford, Wendy Harvey, Tania Carrillo-Roa, Charles P Morris, Tanja Jovanovic, Ross McD Young, Elisabeth B Binder, Joanne Voisey

**Affiliations:** School of Psychology and Counselling, Faculty of Health, Institute of Health and Biomedical Innovation, Queensland University of Technology, Kelvin Grove, Queensland, Australia; School of Biomedical Sciences, Faculty of Health, Institute of Health and Biomedical Innovation, Queensland University of Technology, Kelvin Grove, Queensland, Australia; Gallipoli Medical Research Institute, Greenslopes Private Hospital, Newdegate Street, Greenslopes, QLD 4120, Australia; Department of Translational Research in Psychiatry, Max Planck Institute of Psychiatry, Munich, 80804, Germany; Department of Psychiatry and Behavioral Sciences, Emory University School of Medicine, Atlanta, GA 30322

**Author notes:** Corresponding author Divya Mehta, School of Psychology and Counselling, Faculty of Health, Institute of Health and Biomedical Innovation (IHBI), 60 Musk Avenue, Queensland University of Technology, Kelvin Grove, Queensland, 4059, Australia.

**Keywords:** DNA methylation, biomarkers, combat, stress, PTSD, resilience

## Abstract

Accelerated epigenetic aging, the difference between the DNA methylation-predicted age (DNAm age) and the chronological age, is associated with a myriad of diseases. This study investigates the relationship between epigenetic aging and risk and protective factors of PTSD. Genome-wide DNA methylation analysis was performed in 211 individuals including combat-exposed Australian veterans (discovery cohort, n = 96 males) and trauma-exposed civilian males from the Grady Trauma Project (replication cohort, n = 115 males). Primary measures included the Clinician Administered PTSD Scale for DSM-5 and the Connor-Davidson Resilience Scale (CDRISC). DNAm age prediction was performed using the validated epigenetic clock calculator. Veterans with PTSD had increased PTSD symptom severity (P-value = 3.75 x 10^-34^) and lower CDRISC scores (P-value = 7.5 x 10^-8^) than veterans without PTSD. DNAm age was significantly correlated with the chronological age (P-value = 3.3 x 10^-6^), but DNAm age acceleration was not different between the PTSD and non-PTSD groups (P-value = 0.24). Evaluating potential protective factors, we found that DNAm age acceleration was significantly associated with CDRISC resilience scores in veterans with PTSD, these results remained significant after multiple testing correction (P-value = 0.023; r = 0.32). This finding was also replicated in an independent trauma-exposed civilian cohort (P-value = 0.02; r = 0.23). Post-hoc factor analyses revealed that this association was driven by “self-efficacy” items within the CDRISC (P-value = 0.015). These results suggest that among individuals already suffering from PTSD, some aspects of increased resilience might come at a biological cost.

## Introduction

Posttraumatic stress disorder (PTSD) is a severely debilitating disorder that can develop after experiencing a traumatic event such as sexual assault, natural disaster, life-threatening accidents or combat exposure. Most people experience at least one traumatic event during their lifetime, however only around 14% of those exposed to traumatic events develop PTSD ^1^ A key unresolved question in PTSD research is why certain individuals are at risk of developing PTSD after traumatic exposure, while others appear to be more resilient to the effects of trauma ^2, 3^. Several risk factors for PTSD such as family history of psychiatric disease, childhood trauma, sociodemographic and socioeconomic factors have been established ^4, 5, 6^. In addition, psychological features including hostility, neuroticism and better social functioning have also been identified as predictors of PTSD symptoms ^7, 8, 9^.

How traumatic events are cognitively processed and interpreted influences adjustment and the subsequent development of PTSD ^10, 11^. Allostasis is the active process by which the body responds to daily events and maintain homeostasis while allostatic load or overload is the wear and tear resulting from either too much stress or from inefficient management of allostasis, such as not turning off the stress response when it is no longer needed ^12, 13, 14^ In the context of allostasis, resilience is broadly defined as the ability to adapt successfully in the face of adversity, trauma, tragedy or significant threat ^15, 16^. Individuals who are more resilient are less stressed, less lonely, have better social adaptation skills, and experience greater psychological comfort, therefore resilience has overall positive effects on mental health ^11^. While the majority of research has aimed to identify risk factors for PTSD, fewer studies have specifically looked at resilience in PTSD and how it might moderate the risk of disease. In summary, the neurobiology of risk and resilience in PTSD is dynamic, complex and multilayered with a wide-range of interacting factors.

A major research avenue in the field of PTSD is understanding the biology underlying the disorder. For PTSD, it is clear that both genetic and environmental factors interact with each other to influence disease risk ^17^ The influence of environmental factors on the genome can occur via different means, including epigenetic processes. Epigenetics involves functional alterations in the chromatin structure that can trigger long-lasting modifications in gene expression without creating changes in the DNA sequence ^18^. Among different epigenetic processes such as DNA methylation and histone methylation and acetylation, DNA methylation is one of the common epigenetic mechanism that is involved in physical and mental health and wellbeing ^19, 20, 21^. One aspect of genome-wide epigenetic modifications seen in several disorders, including stress-related disorders, has been demonstrated by the seminal work of Horvath ^22^ who described a robust ‘epigenetic clock’, i.e. a DNA-methylation based predictor of aging. Horvath found a composite predictor comprised of 353 cytosine-phosphate-guanosine sites (CpGs) across the genome (‘epigenetic clock’) that was shown to strongly correlate with chronological age across multiple tissues in humans ^22^, suggesting its usefulness as a biomarker in aging-related research. Using this predictor, accelerated epigenetic aging (Δ - age), defined as the difference between DNA methylation-predicted age (DNAm age) and chronological age, has been associated with aging-related and other phenotypes, including obesity, Down syndrome, Parkinson’s disease, Alzheimer’s and mortality ^23, 24, 25, 26, 27, 28, 29^. We and others recently demonstrated that cumulative lifetime stress accelerated epigenetic aging, an effect that was driven by glucocorticoid-induced epigenetic changes ^30^. Therefore, epigenetic aging is likely to be a key mechanism linking chronic stress with accelerated aging and heightened disease risk for stress-related disorders. To the best of our knowledge, only two studies have tested the association between PTSD and epigenetic aging ^31, 32^, while the role of epigenetic aging in resilience in humans has not yet been studied. In the longitudinal study of Dutch military personnel deployed to Afghanistan, the authors demonstrated that trauma was not associated with decreased telomere length but was associated with accelerated DNAm aging. However, contrary to author expectations, PTSD symptoms were associated with increased telomere length and decreased epigenetic aging ^31^. In the second study, the authors found that DNAm age was not associated with PTSD or neural integrity but accelerated DNAm age was associated with reduced integrity of the corpus callosum genu and poor working memory performance ^32^.

The aim of the present study was to investigate the relationship between epigenetic aging and PTSD and identify underlying risk and potential protective factors.

## Methods

### Samples

The individuals included in the study were part of a larger cohort of Vietnam veterans (N=299) recruited by the Gallipoli Medical Research Foundation (GMRF) at Greenslopes Private Hospital (GPH). Clinical and life experience data, including psychiatric and physical health diagnoses and combat-trauma exposure has been collected for these veterans ^33^. Interview-based data has been supplemented by pre-military and military data from Army records, including information about combat exposure. The study was approved by the Greenslopes Research and Ethics Committee and Queensland University of Technology (QUT) Human Research Ethics Committee and all participants provided written informed consent.

### Clinical Assessments

Structured clinical history included demographics and information regarding smoking, diet and exercise, lifetime history of alcohol consumption, past and current illnesses, medications and family medical history. Structured military and combat history questions such as term of service, number of times served, role and duration in the defence force were also administered for the veterans.

Severity of PTSD symptoms was assessed by clinical psychologists using the Clinician Administered PTSD Scale for DSM-5 (CAPS-5) ^34^ which is a gold-standard for PTSD assessment. Half the veterans had current PTSD symptoms and 25% meet DSM-5 diagnostic criteria for current PTSD. Of this larger sample of 299 veterans, for the current study we selected a sub-sample of veterans with high levels of PTSD symptom severity (n = 48) and veterans with low levels of PTSD symptom severity and no previous PTSD diagnosis (n = 48) as cases and controls respectively, to increase the power to detect biological differences between the groups. Care was taken to match the case-control groups for environmental factors and demographics (Table 1).

Common comorbidities were assessed using the Mini International Neuropsychiatric Interview DSM IV (MINI), an instrument designed to assess major Axis 1 disorders with high validity and reliability ^35, 36^. The Depression Anxiety Stress Scale 21 (DASS-21) is a self-report scale that measures through subscales three different constructs: stress, depression and anxiety ^37^ Higher scores for DASS-21 reflect increased symptoms of depression, anxiety and stress, respectively. The Cronbach’s Alpha DASS-21 was high with an α = 0.95. The Connor-Davidson Resilience Scale (CD RISC) was used to measure resilience via a range of coping strategies that have been shown to be successful mediators in dealing with adversity ^38^. The scale has good psychometric properties, ^39^ with a Cronbach's α = 0.93.

### Experimental Procedures

DNA samples extracted from 8 ml of blood were sent to the Australian Genome Research Facility (AGRF) and stored at -20°C. Quality assessment of the samples was performed by resolution on a 0.8 % agarose gel at 130 V for 60 minutes. Samples were bisulphite converted with Zymo EZ DNA Methylation kit as previously described ^40, 41^. The Infinium platform assays more than 850,000 CpG sites, encompassing 99% of RefSeq genes. It covers 96% of CpG islands with multiple sites in the island, the shores (within 2 kb from CpG islands) and the shelves (>2 kb from CpG islands). It also covers CpG sites outside of CpG islands and DNase hypersensitive sites as well as incorporating miRNA promoter regions. All the Illumina quality controls met performance metrics which included sample-independent, sample dependent, staining, extension, target removal, hybridization, bisulphite conversion I and II, specificity, non-polymorphic and negative controls.

### Replication sample

Replication was performed using the same approach in the Grady Trauma Project (GTP). Details of the cohort are described in Zannas A et al, 2015 ^30^; briefly, the Grady Trauma Project (GTP) is a large study conducted in Atlanta, Georgia, that includes participants from a predominantly African American, urban population of low socioeconomic status ^42, 43^. This population is characterized by high prevalence and severity of trauma over the lifetime and is thus particularly relevant for examining the effects of stressors on epigenetic markers. For replication we used a subset of 115 males that were matched for adult trauma levels (all had two or more types of adult trauma) assessed by the Trauma Events Inventory ^44^ For resilience, like in the discovery sample, the Connor-Davidson Resilience Scale (CD RISC) was used. The GTP samples were run on the Illumina 450k arrays as previously detailed in Zannas et al, 2015 ^30^. Methylation beta values after corrections for batch effects were used for the analysis.

### Statistical analysis

Raw beta values from EPIC Illumina arrays were exported into R for statistical analysis. For quality control (QC), pairs of technical and biological replicates were included to test and confirm the robustness of the microarray procedures. The level of methylation was determined by calculating a “β value” (the ratio of the fluorescent signals for the methylated *vs.* unmethylated sites). Intensity read outs, normalization and methylation beta values calculation were performed using the minfi Bioconductor R package version 1.10.2. For methylation analysis, IDAT files were loaded into the R (2.15) environment using the Bioconductor minfi package (1.4.0) ^45^. The methylation status for each probe was recorded as a *β*-value that ranged between 0 and 1, where values close to 1 represent high levels of methylation and where values close to 0 represent low levels of methylation. A detection P-value was calculated for all probes on all arrays. A *P*-value > 0.05 indicates that the data point is not significantly different from background measurements. Probes with > 50% of the samples with a detection *P*-value > 0.05, probes located on either the X or Y chromosomes and, probes with single-nucleotide polymorphisms present within 10–50 bp from query site, and within < 10 bp from query site were removed. These resulted in a total of *n*=724,232 CpG probes that were used for all subsequent analysis. Data were analysed using an established analysis pipeline comprising of custom statistical programs and scripts ^40, 46, 47^ written in R and Linux. Surrogate variable analyses revealed 17 significant SVA vectors which were used as covariates in the model to correct for technical artefacts and hidden confounds ^48^. DNA methylation-based age prediction was performed using the statistical pipeline developed by Horvath ^22^. For analysis purposes, the DNAm-age was regressed against the chronological age (age acceleration residuals) together with other covariates were used in the generalized regression models. Results were corrected for multiple testing using 10% false discovery rate. Factor analysis of the CD-RISC resilience scores was performed in R using the psych package ^49^. Exploratory analysis using the spree plot and parallel analysis were used to determine the number of significant factors. The minimum residual solution was transformed into an oblique solution using an oblimin transformation and loadings > 0.4 were considered relevant for the factor.

## Results

### Demographics of samples

A total of 96 male Australian veterans from the Vietnam War were included in the study. Demographics and characteristics of the individuals are shown in Table 1. The veterans had an average age of 69 years [SE = 0.45]. A total of 85% veterans were currently married and 79% of the veterans had children. Among the veterans, 8% were working full-time, 10% were working part-time and 66% veterans were currently retired. Veterans with a current diagnosis of PTSD had increased PTSD symptom severity, higher depressive, anxiety and stress scores, increased rates of suicide ideation and significantly higher CDRISC resilience scores compared to veterans without PTSD (Table 1). In the overall sample, PTSD symptom severity was negatively correlated with CDRISC scores (r = -0.58, 5.41 x 10^-10^).

### Relationship between DNA methylation age and PTSD

In the full sample of veterans (n=96), DNAm age was significantly correlated with the chronological age (r = 0.50, P-value = 3.3e-6). No significant differences were present in the DNAm age acceleration between the PTSD and non-PTSD groups (P-value = 0.24). In addition, PTSD symptom severity was not associated with DNAm age acceleration (P-value = 0.47).

### Relationship between DNA methylation age and resilience

Given the significant difference in resilience scores between the PTSD and non-PTSD groups (Table 1), we next sought to investigate the contribution of potential protective factors associated with DNAm age using CDRISC scores. In the overall sample, DNAm age acceleration was not associated with resilience scores (P-value = 0.18). Upon stratification by PTSD diagnosis, we observed that DNAm age acceleration was significantly associated with resilience scores in the PTSD group and remained significant after multiple testing correction (r = 0.32 and P-value = 0.023). No significant correlation was observed in the non-PTSD group (r = -0.19 and P-value = 0.4). Contrary to expectations, we observed that DNAm age acceleration was positively correlated with resilience scores in the PTSD group, such that increased epigenetic age acceleration was associated with increased resilience among veterans with PTSD (Figure 1).

**Figure 1:**
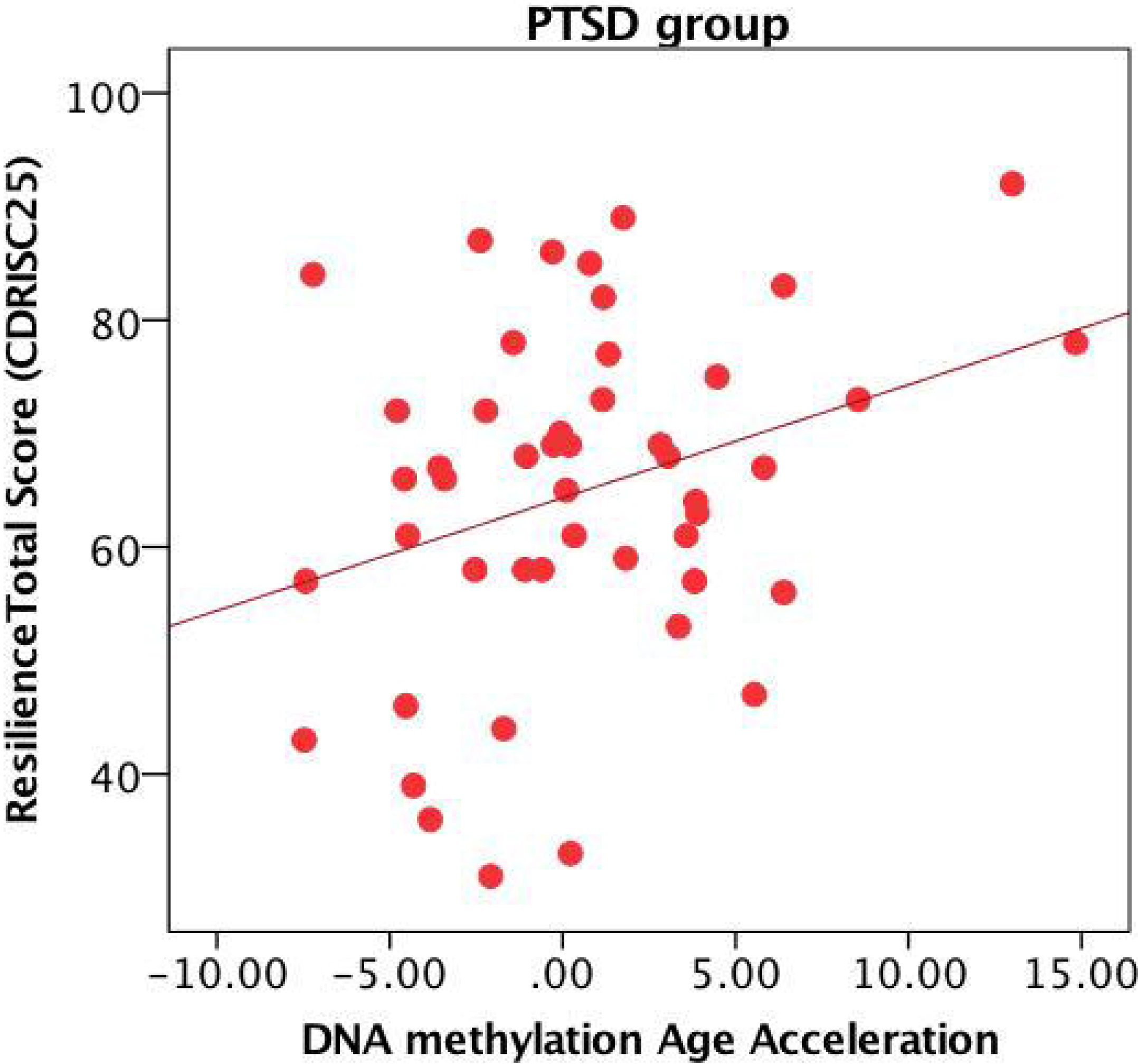
Epigenetic aging and resilience scores: DNAm age acceleration was positively correlated with resilience scores in the PTSD group (r = 0.32 and P-value = 0.023).

Next, we performed post-hoc analyses to investigate if specific latent factors within the Connor-Davidson Resilience Scale might drive the observed positive relationship between DNAm age acceleration and resilience scores in PTSD. Exploratory factor analysis in R using the psych package identified two significant factors, summarizing ‘Hardiness/Adaptability’ and ‘Self-efficacy’ items (Supplementary Figure 1). The 2-factor model fit based upon the diagonal values was 0.96 and a chi-square of 344 with a P-value < 8.2e-05, indicating a good fit. Of the two factors, only the Self-efficacy factor was significantly associated with DNAm age acceleration in the PTSD group (P-value = 0.015).

### Replication of the epigenetic clock and resilience findings in a civilian cohort

We sought to replicate our results in an independent population from the Grady Trauma Project (GTP), which comprises of a population from the suburbs of Atlanta that had been exposed to significant traumatic events ^43^ and test if observed associations between DNA methylation age and resilience scores were specific to the veteran cohort or could be generalized to a non-combat population. We have previously assessed epigenetic aging in this cohort ^30^ and for this replication we used a male subset of the larger sample (n=115). We hypothesized that the relationship between age acceleration and resilience scores would be significantly different between the PTSD and non-PTSD groups.

The replication sample comprised of 115 males (n=70 PTSD and n=45 non-PTSD) aged between 19-65, with a mean age of 44 [1.14]. We tested whether the association between DNAm age acceleration and resilience scores were driven by the PTSD diagnosis status. We observed significant differences in association between age acceleration and resilience scores between PTSD and non-PTSD groups in the GTP cohort (P-value = 0.02). The positive association between age acceleration and resilience scores were also observed in the GTP PTSD group (r = 0.23), similar to that in the discovery sample of veterans (Figure 2). For the non-PTSD group however, in the GTP sample, we observed a stronger negative correlation between age acceleration and resilience (r = -0.25). The results in controls (non-PTSD) from the GTP indicate that as per hypothesis, DNAm age acceleration and resilience are inversely correlated, i.e. increased DNAm age acceleration is associated with decreased resilience. For veterans without PTSD, the same negative correlation between DNAm age and resilience was also observed as described above, but the correlation was slightly blunted (r = -0.19).

**Figure 2:**
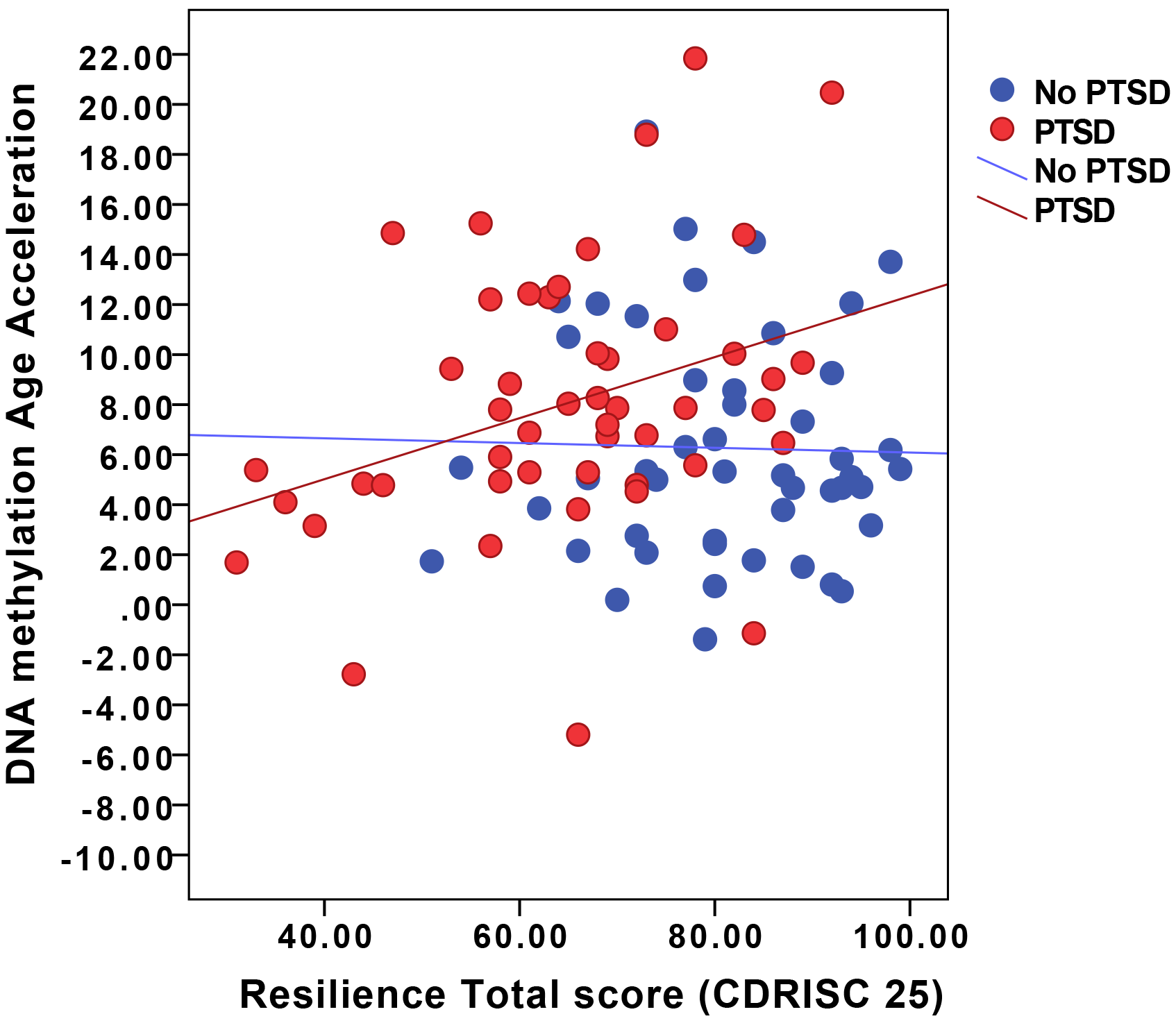
DNAm age acceleration and resilience score – a) DNAm age acceleration was significantly associated with resilience scores (shown in red) in the PTSD group in the Australian Vietnam combat veterans (r = 0.32) and b) DNAm age acceleration was significantly associated with resilience scores (shown in red) in the PTSD group in the replication cohort from the Grady Trauma project (r = 0.23).

These results corroborate our initial findings that increased resilience scores are associated with increased DNAm age acceleration in PTSD and the opposite effect is observed in controls (non-PTSD), with increased resilience scores associated with decreased DNAm age in both the veterans and the civilian population.

## Discussion

In recent years, extensive research effort has been made to identify biological markers that can predict healthy aging as well as disease risk and negative health outcomes associated with aging. In this study, for the first time we assessed the link between the epigenetic clock, a biological marker of aging, and resilience in veterans. The epigenetic clock has been widely described to be associated with detrimental health and a myriad of disorders including obesity, Down syndrome, Parkinson’s disease, Huntington’s disease, and cognitive and physical fitness in the elderly ^23, 24, 25, 26, 27, 28, 29^.

PTSD occurs due to a complex interplay of risk and protective factors that interact with each other to modulate risk of disease. Identification of risk and protective factors for PTSD will help to identify individuals with the greatest necessity for early intervention and facilitate research aimed at improving the efficacy of treatment. In a recent study, higher resilience was associated with improved social functioning after PTSD and depression severity, childhood maltreatment, physical health, gender, education, marital status, and employment were simultaneously adjusted for ^50^. Other studies have illustrated the importance of resilience as a protective factor against PTSD symptoms in high-risk groups ^11^; however, models of resilience include risk as well as protective factors that may interact to reduce negative consequences and facilitate positive consequences ^51^.

Here, we examined resilience measures and demonstrated that while in controls (non-PTSD), increased resilience was associated with decreased DNAm age as expected, among veterans diagnosed with PTSD, increased epigenetic aging was associated with increased resilience, contrary to our expectations. Importantly, in this study, the link between epigenetic aging and resilience among veterans with PTSD was also replicated in an independent sample from a civilian population with high levels of trauma-exposure. In this civilian population, we also observed that increased resilience was associated with increased DNAm age acceleration among individuals with PTSD. These results are in line with previous work showing that elevated allostatic load was observed in women with high stress histories and particularly those with PTSD ^52^. Stress response associated with allostatic load can follow different trajectories including an inability to turn off allostatic responses after stress termination and inadequate responses by some allostatic systems that trigger compensatory increases in other systems ^53, 54, 55^. For example, one study demonstrated that among children in low socioeconomic strata, self-control acts as a “double-edged sword,” facilitating psychosocial adjustment, whilst simultaneously undermining physical health as reflected by increased epigenetic aging ^56^. Our findings also suggest that the allostatic load among individuals with PTSD is distinct from people without PTSD. Among individuals with PTSD, it is likely that there is already an increased allostatic load such that for these individuals, increased resilience further elevates the allostatic load and thereby resulting in an allostatic overload, resulting in a cascading negative impact on overall health.

Factor analysis of the CDRISC scores revealed two significant factors reflecting ‘Adaptability’ and ‘Self-efficacy’, these were similar to other factor analyses of the same resilience inventory ^57, 58, 59^. Of the two factors, we observed that the ‘self-efficacy’ factor was significantly associated with DNAm age acceleration in the PTSD group. Self-efficacy is defined as an individual’s perceived ability to organize and employ one’s skills and execute the necessary actions to achieve desired outcomes ^60, 61^. Previously, Boks and colleagues ^31^ demonstrated that PTSD symptoms were significantly associated with increased telomere length and decreased DNAm aging. The authors discussed differences between short-term and long-term trauma response in aging parameters. The authors noted that their results were consistent with a model whereby aging parameters increase in response to trauma only for those who have already manifested symptoms of PTSD; and for these individuals with PTSD it is likely that this maladaptive state of age decrease subsides afterwards. Wolf and colleagues ^32^ demonstrated that accelerated epigenetic aging may be manifested in degradation in the neural integrity of microstructural cells in the genu in the corpus callosum and working memory deficits and the authors discussed that PTSD may levy a heavy toll on the body reflected, in part, as accelerated cellular aging in the epigenome. Our findings are also in line with previous studies that highlight the notion that self-efficacy is not uniformly beneficial, and that higher levels of self-efficacy can sometimes lead to increases in neuroendocrine and psychological stress responses and decreases in performance, a widely overlooked phenomenon ^62^.

In conclusion, our study sheds light on the role of epigenetic aging in PTSD. This is the first study to evaluate the relationship between epigenetic aging in PTSD by examining the influence of potential protective factors such as resilience. Contrary to our expectations, we found that increased resilience in PTSD is associated with accelerated epigenetic aging and we found that this relationship may be related to underlying self-efficacy. Our findings fit well with the theory of allostatic overload as a result of cumulative physiological wear and tear *via* repeated efforts to adapt to stressors over time. Additional larger, longitudinal studies assessing the effects of stress and epigenetic aging over time would be beneficial to better understand the disease trajectory and biological processes that shape risk and resilience in PTSD.

## Acknowledgements

The authors thank the Gallipoli Medical Research Foundation for their support, especially Miriam Dwyer and Dr Sarah McLeay for their project management support. The authors would also like to acknowledge Dr Madeline Romaniuk for psychological expertise and Dr John Gibson and the team at the Keith Payne Unit, and the staff and investigators at Greenslopes Private Hospital for their valuable contribution to the study. All authors extend their gratitude to the study participants for their generous provision of data and time. The PTSD Initiative was funded by the Queensland Branch of the Returned & Services League of Australia (RSL QLD). The Gallipoli Medical Research Foundation wishes to thank the RSL QLD for their generous donation, and Sullivan Nicolaides Pathology and Queensland X-Ray for their in-kind support. The authors thank QUT and IHBI for financial and administrative support. Divya Mehta was funded via the QUT Vice Chancellors Research Fellowship.

### Potential conflicts of interest

All authors report no potential conflicts of interest.

## Table Legends

Demographics and characteristics of the 96 veterans included in the study.

## Supplementary Information

**Supplementary Figure 1 –.**
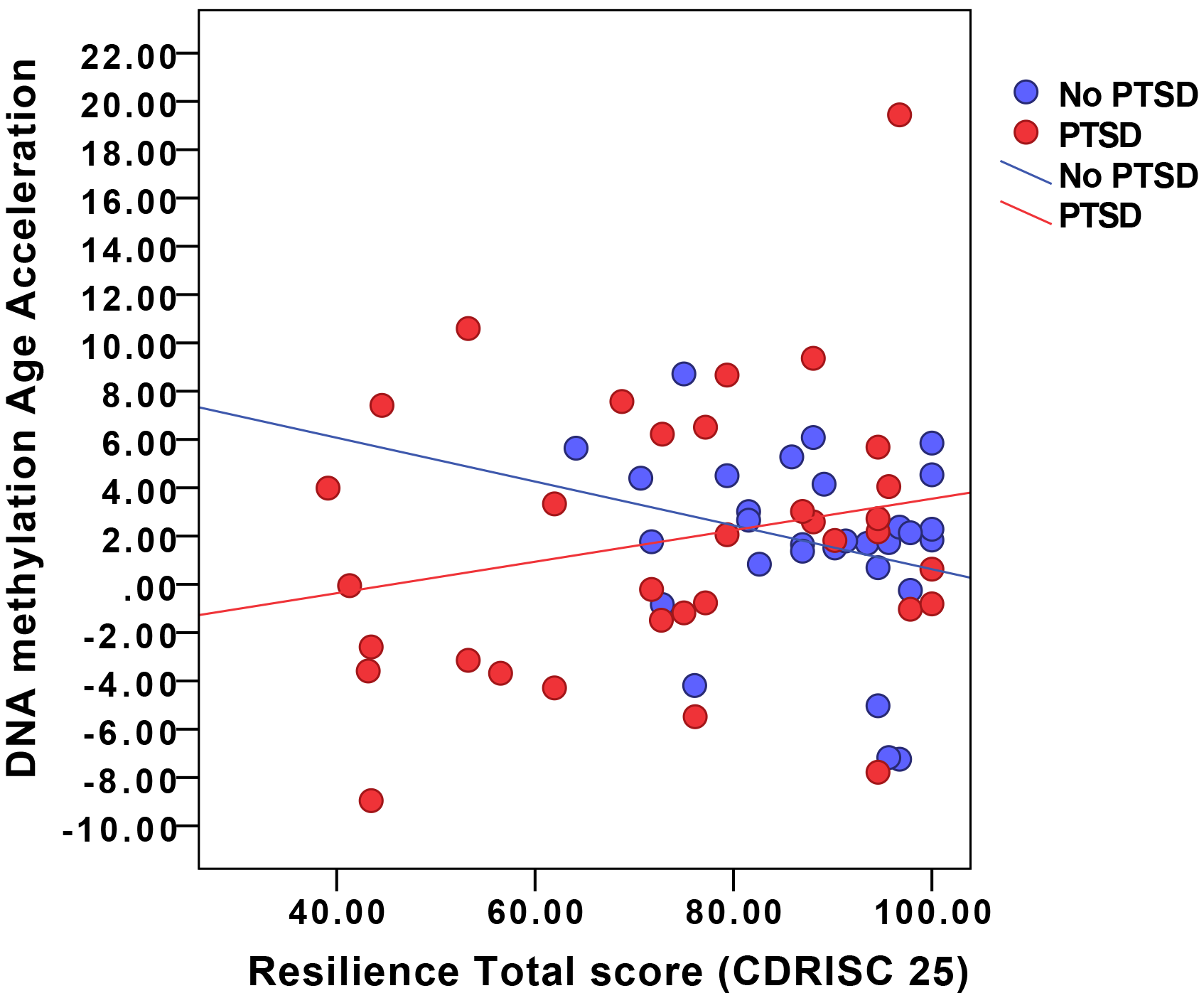
Exploratory factor analysis of the Connor Davidson resilience scores in R using the psych package identified two significant factors. The Cronbach alpha was 0.93, indicating internal consistency and reliability. The 2-factor model fit based upon the diagonal values was 0.96 and the chi-squared of 344 with a P-value <8.2e-05, indicating a good fit.

a. Scree plot and parallel analysis indicating a 2-factor solution is optimal
b. Factor analysis diagram depicting the loadings of the sub-items within the factors. TC1 comprised ‘Hardiness/Adaptability’ and itemsTC2 comprised ‘Self-efficacy’ items

